# Stability and individual variability of social attachment in imprinting

**DOI:** 10.1101/2020.04.04.025072

**Authors:** Bastien S. Lemaire, Daniele Rucco, Mathilde Josserand, Giorgio Vallortigara, Elisabetta Versace

## Abstract

Filial imprinting has become a model for understanding memory, learning and social behaviour in neonate animals. This fast attachment mechanism allows the young of precocial bird species to learn the characteristics of conspicuous visual stimuli and display affiliative response to them. Although more prolonged exposure to an object produces a stronger preference for it afterwards, this relation is not linear. Chicks can even prefer to approach novel rather than familiar objects at some stages of imprinting. The time course and stability of imprinting has just started to be investigated. To date, little is known about how filial preferences develop across time, due to the challenges in assessing individual performance. This study aimed to investigate filial preferences for familiar and novel imprinting objects over time. We have used an automated setup to track the behaviour of chicks continuously for subsequent days. After hatching, chicks were individually placed in an arena where stimuli were displayed on two opposite screens. The duration of exposure and the type of stimuli were manipulated while the time spent at the imprinting stimulus was monitored across six days. We showed that prolonged exposure (3 days vs 1 day) to a stimulus produced robust filial imprinting preferences. Interestingly, with a shorter exposure (1 day), animals re-evaluated their filial preferences in functions of their innate preferences and past experiences. Our study suggests that predispositions influence learning when the imprinting memories are not fully consolidated, driving animal preferences toward more predisposed stimuli.

## Introduction

Young social animals that move around soon after birth, such as ducklings and domestic chicks, require to stay in contact with conspecifics to survive and thrive (Versace & Vallortigara, 2015). It is not surprising, hence, that at the beginning of life they can quickly learn the features of the mother and stay in contact with her, a phenomenon known as filial imprinting (Bateson, 1966; Bolhuis, 1991; Hess, 1959; Lorenz, 1937; McCabe, 2019; Spalding, 1873; Vallortigara & Versace, 2018). In the case of chicks, as little as 15 minutes of visual exposure are sufficient to develop a learned preference for a conspicuous object (Bateson & Jaeckel, 1976). This quickly learned preference has become a model for understanding memory, learning and the onset of social behaviour in neonate animals (Di Giorgio et al., 2017; Rose, 2000, 2003; Solomonia & McCabe, 2015; Versace & Vallortigara, 2015).

Imprinting responses are not only observed in the wild, where they are directed to the mother or siblings (Nicol, 2015). In laboratory settings, chicks imprint on controlled artificial objects (Bolhuis, 1991), such as plastic cylinders (Chiandetti & Vallortigara, 2011; Versace, Schill, Nencini, & Vallortigara, 2016) and computer monitor displays (Santolin, Rosa-Salva, Vallortigara, & Regolin, 2016; Versace, Spierings, Caffini, ten Cate, & Vallortigara, 2017; Wood & Wood, 2015). This paved the way for systematic studies in controlled laboratory conditions.

By exposing chicks to different stimuli, researchers have found that unlearned biases influence filial preferences (Johnson & Horn, 1988; Miura & Matsushima, 2016; Miura, Nishi, & Matsushima, 2020; Versace et al., 2016). Such an effect has been illustrated by Bateson and Jaeckel (1976) by imprinting chicks with a red or yellow object. When the animal received enough exposure to an object, both groups of chicks had a preference for their familiar object. However, the preference was higher in the group of chicks exposed to the red stimulus in comparison to the yellow group. Johnson et al. (1985) found similar results using a red box and/or a stuffed jungle fowl as imprinting and test stimuli. These results suggest that filial preferences are influenced by experience (exposure to an object) and an animal’s predispositions.

Salzen and Meyer (1968) showed that chicks were able to change their imprinting preferences toward a novel object if exposed for a prolonged duration with it. In contrast, other studies (Boakes & Panter, 1985; Bolhuis & Trooster, 1988) showed that imprinting was irreversible if a predisposed stimulus (such as a live hen) was used as primary imprinting stimulus, again suggesting a close relationship between filial preferences and predispositions. Recently, it has been shown that predispositions can affect the acquisition of imprinting memory (Miura & Matsushima, 2016). The authors showed that chicks exposed to a pattern of motion for which they have a spontaneous preference (biological motion) allow them to form a learned colour preference more effectively. Moreover, it has been shown that the combination of predisposed features such as biological motion and red colour located on the chick’s head make imprinting more robust (Miura et al., 2020).

It has been suggested that predispositions direct the attention of the chick toward the kind of stimuli from which the animal would benefit the most (Johnson et al., 1985; Miura & Matsushima, 2016; Miura et al., 2020). In line with this interpretation, chicks have predisposed/not learned preferences for specific patterns of motion (Rosa-Salva, Hernik, Broseghini, & Vallortigara, 2018; Vallortigara, 2012) and arrangments of features (Johnson & Horn, 1988; Rosa-Salva, Mayer, & Vallortigara, 2019; Rosa-Salva, Regolin, & Vallortigara, 2009) that are similar to those found in living animals, such as biological motion (Miura & Matsushima, 2012; Vallortigara, Regolin, & Marconato, 2005), self-propulsion (Rosa-Salva, Grassi, Lorenzi, Regolin, & Vallortigara, 2016; Versace, Ragusa, & Vallortigara, 2019) or even specific colours such as red (which is the colour of the comb, a specific zone of the head that is known to convey important physiological information, Guhl & Ortman, 1953).

Colours are crucial to discriminate between individuals in a chicken flock (Guhl & Ortman, 1953). In filial imprinting, it has also been described to be an essential characteristic used by the young animal to recognize their artificial objects (Maekawa et al., 2006). Some colours are more effective than others in imprinting (Bolhuis, 1991). Although the effect of the contrast between a colour and its background has not been clarified yet, red, orange and blue appear to elicit stronger responses than green and yellow (Ham & Osorio, 2007; Kovach, 1971; Salzen, Lily, & McKeown, 1971; Schaefer & Hess, 2010). Therefore, red and blue can be considered as “predisposed” imprinting stimuli. While spontaneous preferences have been described for specific colours, whether those preferences are steady or can change in time has not been investigated.

Filial imprinting preferences have been well described (Bolhuis, 1991; McCabe, 2019). However, how these preferences develop in time and vary depending on the imprinting objects used (more or less attractive) and long duration of imprinting has not been documented. From this extensive literature, we know that longer exposure produces stronger preferences for the imprinting stimulus (familiar stimulus) (Bateson & Jaeckel, 1976; Bolhuis, Cook, & Horn, 2000; Hess, 1959). However, after imprinting the preference for approaching familiar objects and avoiding novel objects is not merely steady nor incremental. On the contrary, it has been repeatedly shown that in some situations, chicks prefer to approach novel rather than familiar objects, an unexpected behaviour. For instance, Bateson and colleagues (Bateson & Jaeckel, 1976; Jackson & Bateson, 1974) have observed that in the initial stage of imprinting – i.e. 15 and 30 minutes after the beginning of the imprinting phase but not after 60 minutes – chicks are motivated to be exposed to novel objects. In this setting, chicks actively actioned a lever to be exposed to an alternative object (Jackson & Bateson, 1974). More recently, the early shift from the first object to the exploration of alternative stimuli has been observed in different breeds of chicks that were tested on their spontaneous preferences to approach a stuffed hen versus a scrambled version of it. Versace et al. (2017) have shown that while in the first 5 minutes of visual experience all breeds had a preference for the stuffed hen, 5 minutes later two breeds started to explore also the other stimulus. Interestingly, preferences for novel stimuli were also documented to appear much later using imprinting paradigms (Versace, Regolin, & Vallortigara, 2006; Versace, Spierings, et al., 2017).

While it has been shown that exploration of novelty takes place at different stages of imprinting, how and why this counterintuitive phenomenon appears remains an open question. To date, the transient preference for unfamiliar stimuli, named ‘slight-novelty preference’ by Bateson (1973), has been described and modelled as an exploration of different points of view of the imprinting stimulus. According to this hypothesis, the preference for exploring objects slightly different from the imprinting stimulus would be useful to recognise different points of view of the mother hen and build a complete representation of it (Bateson, 1979; Jackson & Bateson, 1974; McCabe, 2019). This hypothesis is supported by other studies showing that when two stimuli are presented in close temporality, they became “blended” as a unique stimulus for the animals (Chantrey, 1974; Honey & Bateson, 1996). It has been documented that chicks were learning slower to distinguish two stimuli previously displayed in fast alternation in comparison to slow (Chantrey, 1974). However, the hypothesis that (only) in the first hour of exposure chicks explore novel stimuli to improve the representation of the imprinting object has been confuted. Chicks are still consolidating the representation of the imprinting object much after the first 60 minutes of exposure, as shown by the fact that biochemical learning-related changes are still observed more than 15 hours after the start of imprinting and affected by sleep (Jackson et al., 2008; Solomonia et al., 2003; Solomonia et al., 2011; Solomonia & McCabe, 2015).

There are examples in which novelty preference has been observed much after the early stages of imprinting. On day four post-hatching, after three days of imprinting, chicks had a preference for novel patterns of grammar (Versace et al., 2006). A similar result had been described more recently while investigating a spontaneous generalization of abstract multimodal pattern in chicks (Versace, Spierings, et al., 2017). Interestingly, the preferences were different between sexes. Males showed a preference for unfamiliar stimuli, whereas females showed a preference for familiar stimuli. These results are consistent with sex differences described until know in the context of social recognition. It has been described that females spend more time close to familiar individuals, whereas males prefer to spend more time close to unfamiliar individuals (Vallortigara & Andrew, 1991; Vallortigara, 1992).

Besides the variables influencing filial preferences, chicks might exhibit consistent individual differences. Due to technological limitations, little has been investigated yet. Templeton and Smith (1966) described that chicks response to an effective stimulus were varying across a wide range of performance and was not affected by genetics. More recently, Gribosvkiy and collaborators (Gribovskiy et al., 2015) developed a quantitative methodology to study the inter-individual variability among chicks in imprinting and showed the existence of high variability between individuals. Moreover, individual variability is influenced by different behavioural types. For example, chicks with higher behavioural flexibility have been described to have a stronger preference for novelty in a generalization task (Zidar, Balogh, Leimar, & Løvlie, 2019). Thanks to recent progress in technology and tracking techniques, it is now possible to investigate animals behaviours in time reliably (Anderson & Perona, 2014; Goldman & Wood, 2015; Nath et al., 2019; Versace, Caffini, Werkhoven, & de Bivort, 2020). Therefore, offering the possibility to study individual preferences across consecutive days automatically.

In this study, we have built an automated setup where we continuously tracked the behaviour of chicks from the first exposure to the imprinting stimuli for six subsequent/consecutive days. Chicks were individually housed in an arena with two opposite monitors. The position of imprinting and test stimuli was counterbalanced between monitors while we kept track of the distance of chicks from the stimuli. Imprinting duration and testing duration were manipulated. Objects of different colours were used as imprinting objects to investigate its effect on chicks preferences between the imprinting object and an unfamiliar one. In Experiment 1, chicks were imprinted for 1 day with one stimulus and tested for 5 days with two stimuli. In Experiment 2, the imprinting duration was increased to 3 days and chicks were tested for 3 days too. In Experiment 3, chicks were imprinted with one object for 1 day, then with another one for 2 days and tested for 3 days. In Experiment 4, we replicated a similar procedure than Experiment 3, but this time assessed the animal preference between their primary or secondary imprinting object. In such settings with prolonged and continuous behavioural monitoring, we investigated how filial preferences developed in time and looked at individual preferences.

## Materials and Methods

### Subjects

We used 128 domestic chicks (*Gallus gallus*) of the strain Ross 308 (a strain selected to be sexually dimorphic at birth, based on the feathers). This study was carried out in compliance with the European Union and the Italian law on the treatment of animals. The experimental procedures were approved by the Ethical Committee of the University of Trento and licenced by the Italian Health Ministry (permit number 53/2020). The eggs were coming from a commercial hatchery (Azienda Agricola Crescenti) and were incubated at the University of Trento under standards controlled conditions (37.7°C and 40% of humidity). Three days before hatching eggs were transferred into opaque individual boxes within a hatching chamber (37.7°C and 60% of humidity).

### Setup

Several apparatuses were used simultaneously. Each apparatus had a rectangular shape (90 cm x 60 cm x 60 cm, Figure 1). A high-frequency computer screen (ASUS MG248QR, 120 Hz) was located on each smaller wall and used to display stimuli. A Microsoft life camera was located on the top of the apparatus at 105 cm from the ground to record the behaviours of the animal. Food and water were located in the middle of the apparatus and available ad libitum.

**Figure 1:**
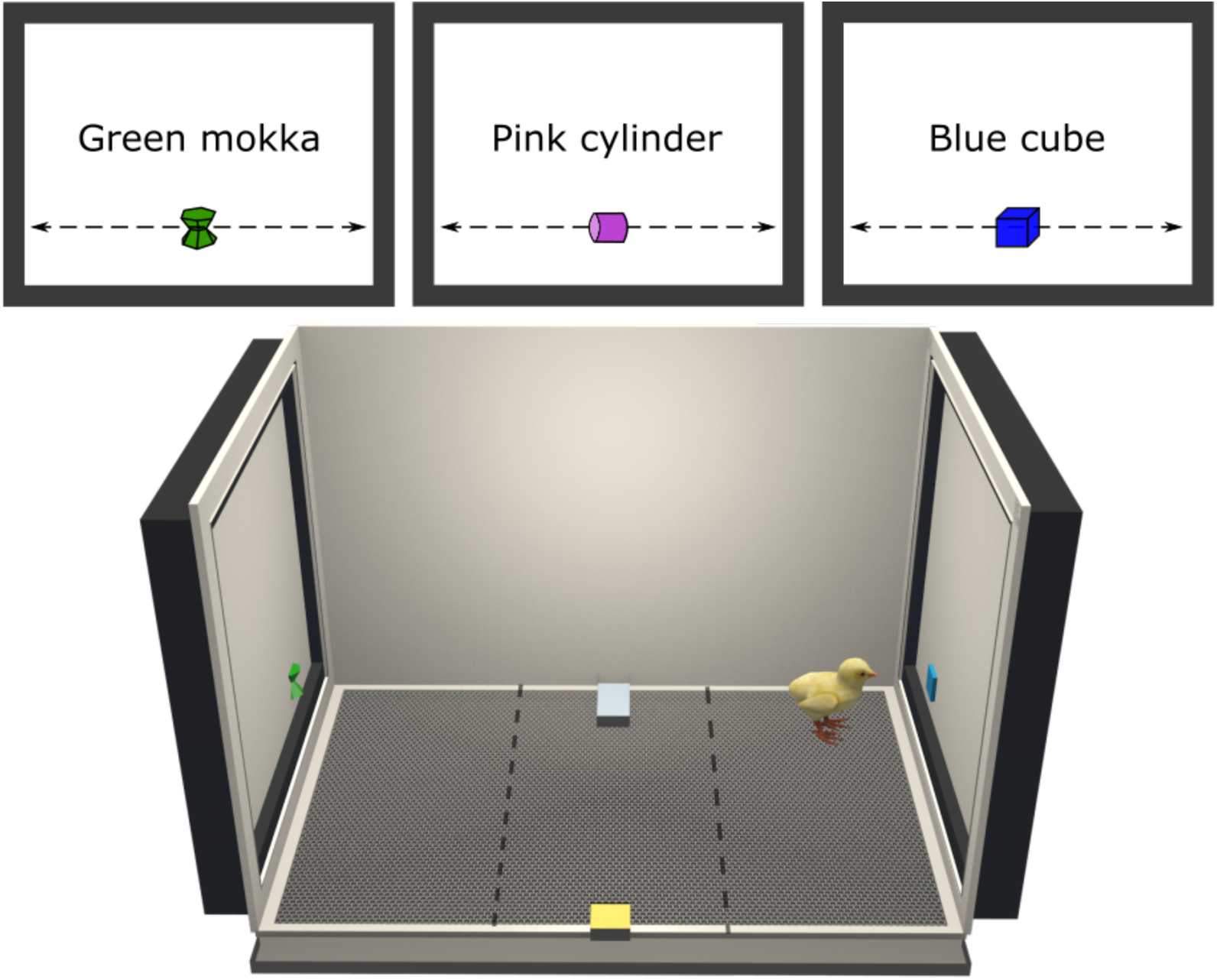
Three-dimensional representation of the apparatus and stimuli used in this study. The stimuli were moving horizontally alongside the screens to attract the attention of the animals. The filial preference of a chick was revealed by its choice to remain near the stimuli displayed. The dashed lines show the delimitation of the virtual zones used to assess the preference of the animal. The time spent near the stimuli was monitored to calculate a Preference for the imprintings stimuli.

### Stimuli

Three-dimensional virtual visual stimuli were created (Figure 1) and animated on Blender (v2.79). The objects were different in term of colours and shapes but had similar sizes (5 cm x 5 cm, figure 1). The stimuli were animated (linear movement) in a 3D environment and were crossing the screen in 4.5 seconds (from left to right, see videos in supplementary materials). The video displaying the stimuli was exported with a high frame frequency (120 frames per second, fps).

### General procedure

After hatching, chicks were sexed and individually placed in their apparatus for six days in a day-night cycle (14:10 hours). During the day, the chicks were exposed to the stimuli displayed on the screens. The displaying of the stimuli was divided into different sessions depending on the experimental phase (form 2 hours to 30 minutes). The position of the stimuli on the screens was counterbalanced across sessions. During the night, dark screens were displayed.

Four different experiments were performed. Each experiment was divided into 2 or 3 different phases (primary imprinting, secondary imprinting and testing) and conditions (blue and green). The duration of each phase was manipulated from one experiment to another.

#### Primary Imprinting

This phase was the first one of each experiment. The chicks were exposed to a single imprinting stimulus (the blue or the green depending on the condition). The imprinting sessions lasted two hours (7 sessions) on the first primary imprinting day and one hour on the following days (13 sessions interrupted by 5 minutes period of dark screens).

#### Secondary imprinting

In Experiment 3 and 4, this phase followed the primary imprinting phase and lasted 2 days. The chicks were exposed to a new stimulus (a pink cylinder). The sessions were lasting one hour (13 sessions interrupted by 5 minutes period of dark screens).

#### Testing

Depending on the experiment, the testing phase was either following the primary (Experiment 1 and 2) or secondary imprinting phase (Experiment 3 and 4). The chicks were exposed to two stimuli (primary imprinting stimulus vs novel stimulus or primary vs secondary imprinting stimulus), and their preferences were monitored. The sessions lasted thirty minutes (24 sessions interrupted by 5 minutes period of dark screens between each session).

### Experiment 1

Chicks were exposed to an imprinting stimulus for 1 day (blue or green stimulus depending on the condition) and then tested with two stimuli (imprinting stimulus vs unfamiliar stimulus) for 5 days (figure 2A).

**Figure 2:**
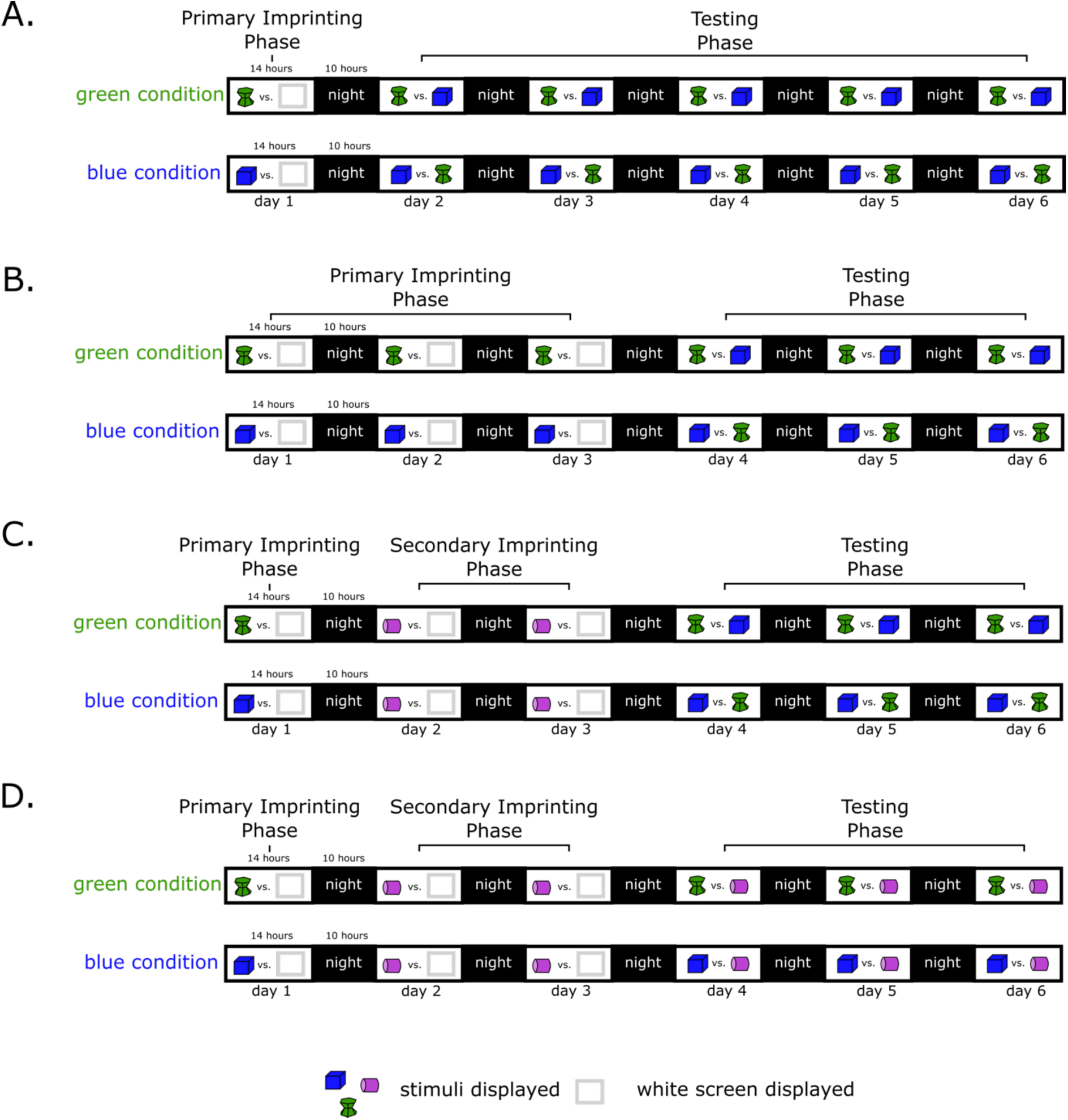
Experimental timelines of experiment 1 (A.), 2 (B.), 3 (C.) and 4 (D.).

Subjects: We imprinted 16 animals (8 females, 8 males) with the green stimulus (green condition) and 16 animals (8 females, 8 males) with the blue cube (blue condition).

### Experiment 2

Chicks were exposed to an imprinting stimulus for 3 days (blue or green stimulus depending on the condition) and then tested with two stimuli (imprinting stimulus vs unfamiliar stimulus) for 3 days (figure 2B).

Subjects: We imprinted 16 animals (8 females, 8 males) with the green stimulus (green condition) and 16 animals (8 females, 8 males) with the blue cube (blue condition).

### Experiment 3

Chicks were exposed to a primary imprinting stimulus for 1 day (blue or green stimulus depending on the condition), secondary imprinting stimulus (pink stimulus) for 2 days and then tested with two stimuli (primary imprinting stimulus vs unfamiliar stimulus) for 3 days (figure 2C).

Subjects: We imprinted 16 animals (8 females, 8 males) with the green stimulus (green condition) and 16 animals (8 females, 8 males) with the blue cube (blue condition).

### Experiment 4

Chicks were exposed to a primary imprinting stimulus for 1 day (blue or green stimulus depending on the condition), secondary imprinting stimulus (pink stimulus) for 2 days and then tested with two stimuli (primary imprinting stimulus vs secondary imprinting stimulus) for 3 days (figure 2D).

Subjects: We imprinted 16 animals (8 females, 8 males) with the green stimulus (green condition) and 17 animals (8 females, 9 males) with the blue cube (blue condition).

### Data analysis

The position of the animal was analyzed automatically using DeepLabCut, an open-source deep-learning toolbox made to track efficiently animal behaviours (Nath et al., 2019). The preference for a stimulus was assessed using the time spent inside the closest zone to it (30 cm wide). The apparatus had been virtually divided into three equal zones corresponding to the left, centre and right side of each arena (Figure 1).

#### Imprinting phases

During these phases (primary and secondary), the number of seconds [s] spent close to the stimulus (in the 30 cm zone close to the screen) was analysed to check for the amount of time spent attending the imprinting object.

#### Testing phase

For this phase, the Preference for the imprinting stimulus [%] was calculated using the following formula:

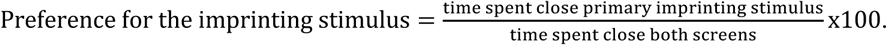

Using this formula, a score of 50 % indicates no preference for either stimulus. A score higher than 50% indicates more time spent at the primary imprinting object. A score lower than 50 % indicates more time spent at the unfamiliar stimulus (Experiment 1, 2 and 3) or the secondary imprinting object (Experiment 4).

### Statistical analysis

#### Imprinting phases

To assess the time spent by the chicks close to the imprinting stimulus during the imprinting phases (primary and secondary), we used an ANOVA with seconds spent close to the imprinting stimulus as dependent variable and Condition (imprinted with green, imprinted with blue), Sex (female, male). In all experiments, data met assumptions of parametric analyses.

#### Testing phase

To determine whether chicks had different preferences for the imprinting stimulus (or the primary imprinting stimulus) between Condition (imprinted with green, imprinted with blue), Sex (female, male) and Day (experiment 1: day 2, 3, 4, 5, 6; other experiments: day 4, 5, 6), we performed a mixed-design ANOVA for each testing phase. To meet parametric analysis assumptions, we arcsin transformed the data. To check whether chicks had a significant preference for the imprinting stimulus or unfamiliar stimulus (primary vs secondary imprinting stimulus in experiment 4) we performed two-tailed one-sample t-tests vs the chance level (50%). Since the chicks underwent several imprinting and testing sessions across testing days, it was possible to test their preference individually. Individual preferences were assessed and compared from chance-level (50%) using two-tailed one-sample t-tests. In each experiment, Levene’s test was conducted to explore chicks variability between conditions (imprinted with green or imprinted with blue). For all experiments, we used an α = 0.05. Analyses were performed using RStudio v1.1 (RStudio Team, 2015). The following packages were used: *goftest* (Faraway, Marsaglia, Marsaglia, & Baddeley, 2019), *nlme* (Pinheiro, Bates, DebRoy, Sarkar, & R Core, 2020), *lme* (Bates, Mächler, Bolker, & Walker, 2015), *tidyr* (Wickham & Lionel, 2020), *plyr* (Wickham, 2011), *dplyr* (Wickham, François, Henry, & Kirill, 2020), *reshape* (Wickham, 2007), *lsr* (Navarro, 2015), *ggplot2* (Wickham, 2016).

## Results

### Experiment 1

#### Imprinting

There were non-significant effects of Condition (*F*(1, 28) = 0.57, *p* = 0.46), Sex (*F*(1, 28) = 0.18, *p* = 0.67) or interaction Sex x Condition, *F*(1, 28) = 0.14, *p* = 0.71) on the time spent close to the imprinting stimulus. The chicks significantly remained closer to the imprinting stimulus (*t*(31) = 83.25, *p* < 0.001, Cohen’s *d* = 14.72), spending 96% of their time (+/- 0.56 SEM) close to the imprinting stimulus. Individual preferences were calculated and showed that all 32 chicks remained significantly more on the side of the arena in which the imprinting stimulus was displayed (Table 1).

#### Testing

The results are shown in *Figure 3*. There were non-significant effects of Condition (*F*(1, 28) = 0.89, *p* = 0.37), Sex (*F*(1, 28) = 0.50, *p* = 0.49), Day (*F*(4, 112) = 1.06, *p* = 0.38) or interactions (Sex x Condition *F*(1, 28) = 0.009, *p* = 0.93; Sex x Day, *F*(4, 112) = 0.28, *p* = 0.89; Sex x Condition x Day, *F*(4, 112) = 0.40, *p =* 0.81), but a significant interaction between Day and Condition on the Preference for the imprinting stimulus (*F*(4, 112) = 2.69, *p* < 0.05). Post hoc analysis (Tukey) showed that the preference for the imprinting stimulus observed on day 2 was significantly different from the preference observed on day 4 in the green condition (*t*(112) = 3.52, *p* < 0.05, Cohen’s *d* = 0.74). On day 2, chicks had a significant preference for the imprinting stimulus (*t*(15) = 4.45, *p* < 0.001, Cohen’s *d* = 1.12) and spent 65% (+/- 3.31 SEM) of their time close to it. However, on day 4, chicks had no preference (*t*(15) = 0.33, *p* = 0.75, Cohen’s *d* = 0.082) and spent 52% (+/- 5.26 SEM) of their time close to their imprinting stimulus. The post hoc test did not reveal other differences. Chicks imprinted with the blue stimulus had a significant and stable preference for the imprinting stimulus (*t*(15) = 3.83, *p* < 0.01, Cohen’s *d* = 0.96) and spent 62% (+/- 3.23 SEM) of their time close to it.

**Figure 3:**
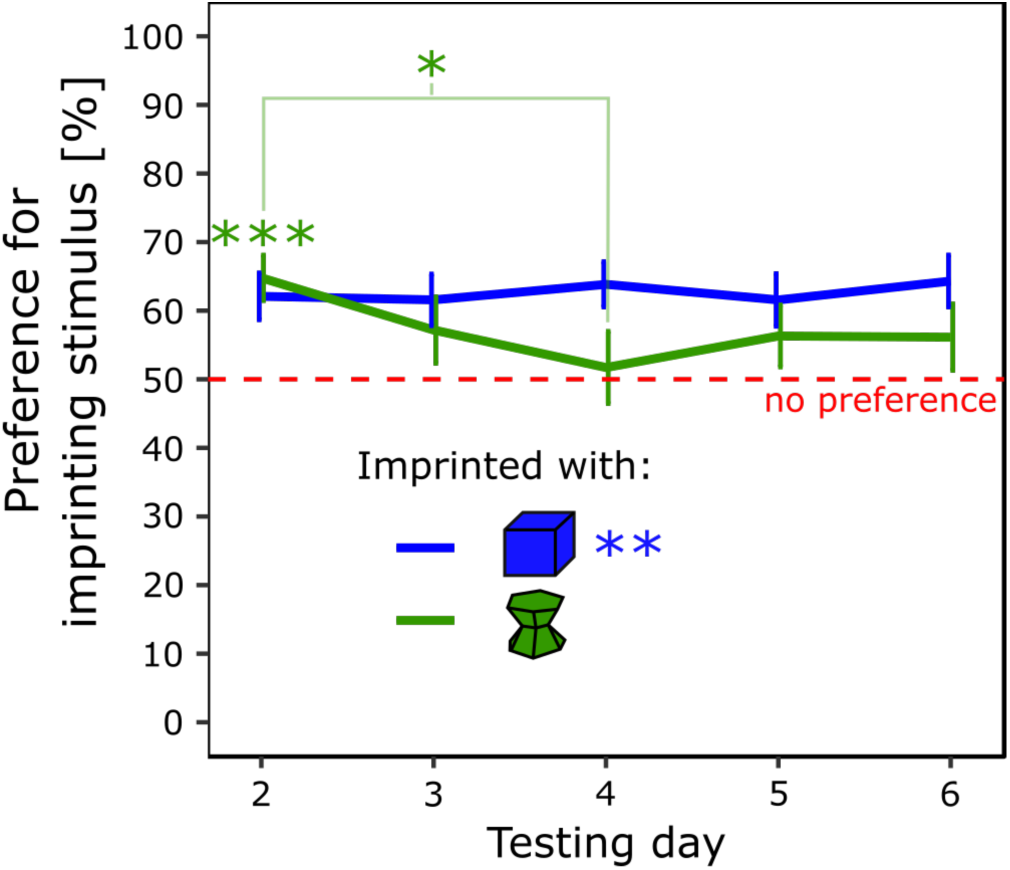
Preference for the imprinting stimulus for each testing day and condition (p < 0.05, *; p < 0.01, **; p < 0.001, ***). The blue line represents the Preference score of the chicks imprinted with the blue stimulus. The green line represents the Preference score of the chicks imprinted with the green stimulus.

Individual preferences were calculated. In the blue condition, 10 chicks (63%) had a significant preference for the imprinting stimulus, 5 (31%) had no preference, and 1 (6%) significantly preferred the unfamiliar stimulus. In the green condition, 7 chicks (44%) had a significant preference for the imprinting stimulus, 6 (37%) had no preference, and 3 (19%) had a significant preference for the unfamiliar stimulus (Table 1 in the supplementary material). Levene’s test showed that the variances of the two conditions were similar (*F*(1, 30) = 0.32, *p* = 0.86).

### Experiment 2

#### Imprinting

There were non-significant effects of Condition (*F*(1, 28) = 1.15, *p* = 0.29), Sex (*F*(1, 28) = 0.002, *p* = 0.97) or interaction (Sex x Condition, *F*(1, 28) = 3.3, *p* = 0.08) on the time spent close to the imprinting stimulus. The trend revealed above was induced by an opposite pattern of between males and females within each condition with small variances. Nonetheless, the time spent close to the imprinting stimulus between each group was similar. Overall, the chicks significantly remained close the imprinting stimulus (*t*(31) = 49.92, *p* < 0.001, Cohen’s *d* = 8.82) 93% of their time (+/- 0.46 SEM).

Individual preferences were calculated and showed that all 32 chicks chose significantly more the side of the arena, where the imprinting stimulus was displayed (Table 2 in the supplementary material).

#### Testing

The results are shown in *Figure 4*. There were non-significant effects of Condition (F(1, 28) = 2.90, p = 0.10), Sex (F(1, 28) = 2.12, p = 0.16), Day (F(2, 56) = 0.63, p = 0.54) or interactions (Sex x Condition, *F*(1, 28) = 0.003, *p* = 1.0; Sex x Day, *F*(2, 56) = 0.05, *p* = 0.95, Condition x Day, *F*(2, 56) = 0.46, *p* = 0.63; Sex x Condition x Day, *F*(2, 56) = 1.52, *p* = 0.23) on the Preference for the imprinting stimulus. The preference for the imprinting stimulus was significantly different from chance-level (*t*(31) = 6.58, *p* < 0.001, Cohen’s *d* = 1.16). The chicks spent on average 69% (+/- 2.90 SEM) of their time close to their imprinting stimulus.

**Figure 4:**
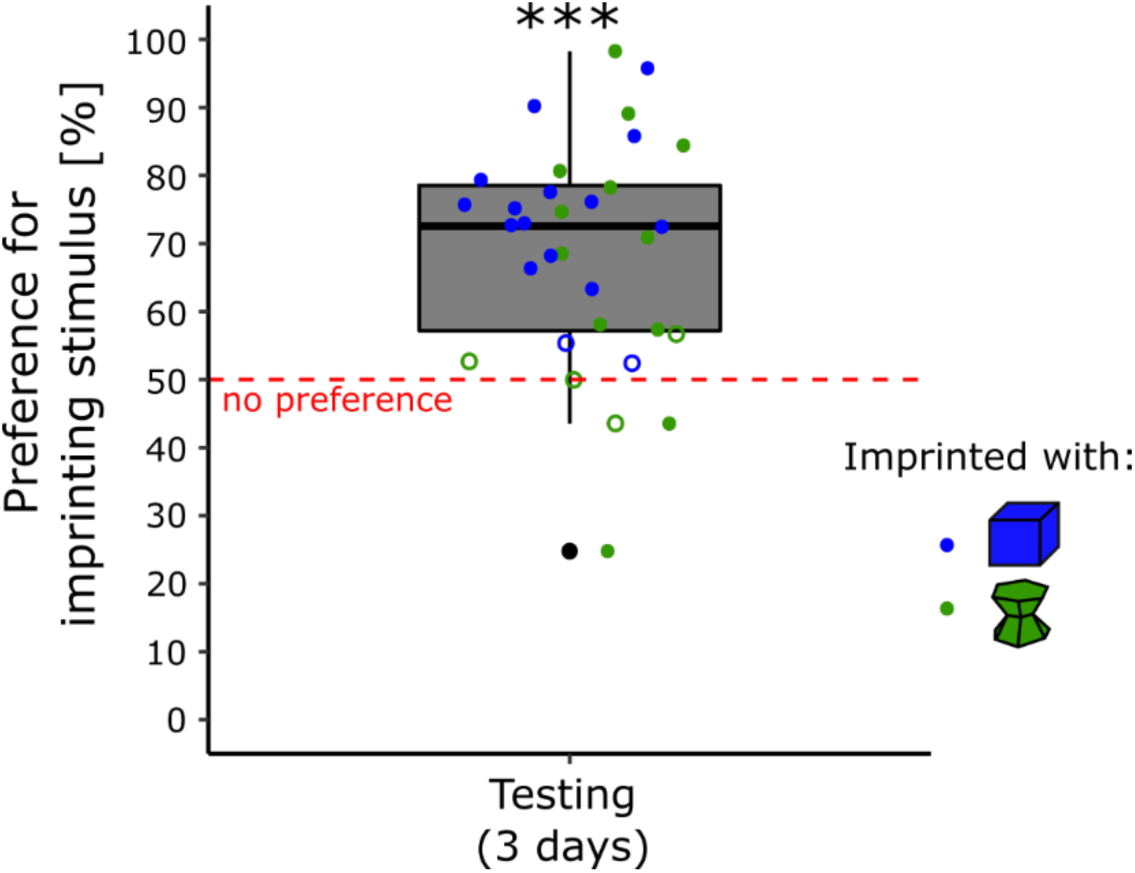
Preference for the imprinting stimulus during the testing phase (p < 0.001, ***). The blue dots represent the preference score of the chicks imprinted with the blue stimulus. The green dots represent the preference score of the chicks imprinted with the green stimulus. Filled dots show the individuals having a significant preference while empty dots show the individuals having no preference.

Individual preferences were calculated. In the blue condition, 14 chicks (87.5%) had a significant preference for the imprinting stimulus, 2 (12.5%) had no preference, and none (0%) significantly preferred the unfamiliar stimulus. In the green condition, 10 chicks (62.5%) had a significant preference for the imprinting stimulus, 4 (25%) had no preference, and 2 (12.5%) had a significant preference for the unfamiliar stimulus (Table 2 in the supplementary material). Levene’s test showed that the variances of the two conditions were significantly different (*F*(1, 30) = 6.14, *p* < 0.05). Chicks imprinted with the green stimulus showed higher variability in their preferences for the imprinting stimulus during testing (s2 = 380.85) than chicks imprinted with the blue stimulus (s2 = 129.91).

### Experiment 3

#### Primary imprinting

There were non-significant effects of Condition (*F*(1, 29) = 0.52, *p* = 0.48), Sex (*F*(1, 29) = 0.17, *p* = 0.69) or interaction (Sex x Condition, *F*(1, 29) = 1.62, *p* = 0.21) on the time spent close to the primary imprinting stimulus. The chicks significantly remained close the primary imprinting stimulus (*t*(32) = 87.18, *p* < 0.001, Cohen’s *d* = 15.18) 97% of their time (+/- 0.54 SEM).

Individual preferences were calculated and showed that all 33 chicks remained significantly more on the side of the arena, where the primary imprinting stimulus was displayed (Table 3 in the supplementary material).

#### Secondary imprinting

There were non-significant effects of Condition on the time spent close to the secondary imprinting stimulus (*F*(1, 29) = 0.14, *p* = 0.72), Sex (*F*(1, 29) = 0.49, *p* = 0.49) or interaction (Sex x Condition, *F*(1, 29) = 0.70, *p* = 0.41) on the time spent close to the secondary imprinting stimulus. The chicks significantly remained close the secondary imprinting stimulus (*t*(32) = 34.72, *p* < 0.001, Cohen’s *d* = 6.04) 93% of their time (+/- 1.25 SEM).

Individual preferences were calculated and showed that all 33 chicks remained significantly more on the side of the arena, where the secondary imprinting stimulus was displayed (Table 3 in the supplementary material).

#### Testing

The results are shown in *Figure 5*. There was a significant effect of Condition (*F*(1, 29) = 70.35, *p* < 0.001) but non-significant effects of Sex (*F*(1, 28) = 2.98, *p* = 0.095), Day (*F*(2, 58) = 0.54, *p* = 0.59) or interactions (Sex x Condition, *F*(1, 29) = 1.21, *p* = 0.28; Sex x Day, *F*(2, 58) = 0.072, *p* = 0.93, Condition x Day, *F*(2, 58) = 0.41, *p* = 0.67; Sex x Condition x Day, *F*(2, 58) = 0.010, *p* = 0.10) on the preference for the primary imprinting stimulus

**Figure 5:**
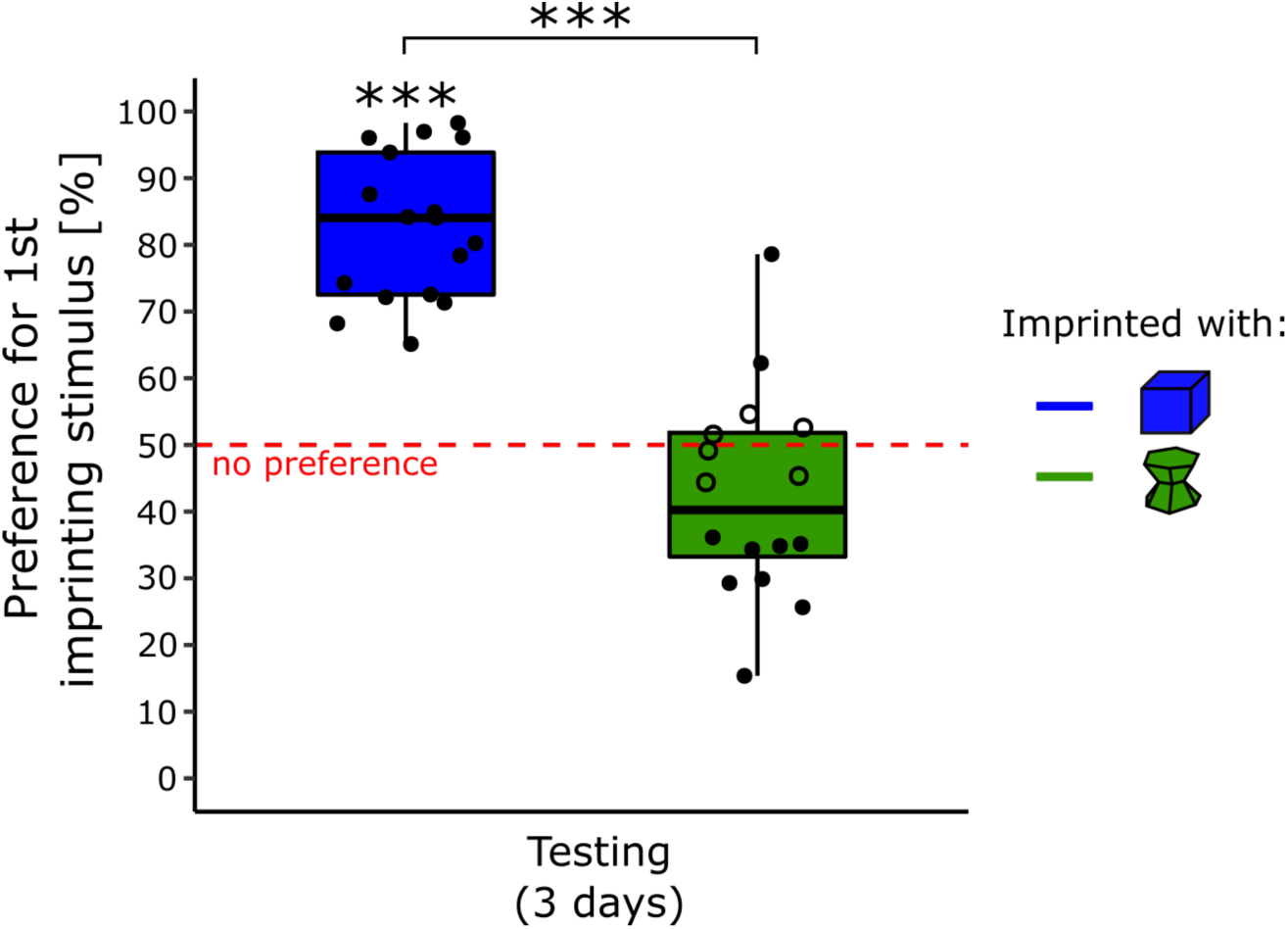
Preference for the primary imprinting stimulus for each condition during the testing phase (p < 0.001, ***). The blue line represents the preference score of the chicks imprinted with the blue stimulus. The green line represents the preference score of the chicks imprinted with the green stimulus. Filled dots show the individuals having a significant preference while empty dots show the individuals having no preference.

The preference for the primary imprinting stimulus was significantly different from chance level for the chicks imprinted with the blue stimulus (*t*(16) = 12.27, *p* < 0.001, Cohen’s *d* = 2.98, *Bonferroni* correction) with an average time spent close to the primary imprinting stimulus of 83 % (+/- 2.66 SEM). The Preference score was non-significantly different from chance level for the chicks imprinted with the green stimulus (*t*(15) = -1.94, *p* = 0.14, Cohen’s *d* = 0.48, Bonferroni correction) with an average time spent close to the primary imprinting stimulus of 42 % (+/- 3.90 SEM).

Individual preferences were calculated and showed that all the chicks (17) had a significant preference for the imprinting stimulus while primary imprinted with the blue stimulus (Table 3 in the supplementary material). Whereas for the chicks primarily imprinted with the green stimulus, 2 (13%) had a significant preference for their primary imprinting stimulus, 6 (37%) had no preference and 8 (50%) had a preference for the unfamiliar stimulus (Table 3 in the supplementary material). Levene’s test showed that the variances of the two conditions were similar (*F*(1, 31) = 1.45, *p* = 0.24).

### Experiment 4

#### Primary imprinting

There were non-significant effects Condition (*F*(1, 29) = 3.44, *p* = 0.074), Sex, (*F*(1, 29) = 0.50, *p* = 0.23) or interaction (Sex x Condition, *F*(1, 29) = 0.10, *p* = 0.75) on the time spent close to the primary imprinting stimulus. The chicks significantly remained close the primary imprinting stimulus (*t*(32) = 45.53, *p* < 0.001, Cohen’s *d* = 7.93) 95% of their time (+/- 0.99 SEM).

Individual preferences were calculated and showed that 32 (97%) chicks remained significantly more on the side of the arena, where the primary imprinting stimulus was displayed, and 1 (3%) did not (Table 4 in the supplementary material).

#### Secondary imprinting

There were non-significant effects of Condition on the time spent close to the secondary imprinting stimulus (*F*(1, 29) = 0.27, *p* = 0.61), Sex (*F*(1, 29) = 0.002, *p* = 0.96) or interaction (Sex x Condition, *F*(1, 29) = 0.30, *p* = 0.59) on the time spent close to the secondary imprinting stimulus. The chicks significantly remained close the secondary imprinting stimulus (*t*(32) = 40.27, *p* < 0.001, Cohen’s *d* = 7.01) 93% of their time (+/- 1.07 SEM).

Individual preferences were calculated and showed that all 33 chicks chose significantly more the side of the arena where the secondary imprinting stimulus was displayed (Table 4 in the supplementary material).

#### Testing

Two chicks (2 males of the blue condition) were removed from the following analyses because the video recordings of their last testing day went missing (camera crash). The results are shown in *Figure 6*. There were non-significant effects of Condition (*F*(1, 27) = 0.11, *p* = 74), Sex (*F*(1, 27) = 2.22, *p* = 0.15), Day (*F*(2, 54) = 0.14, *p* = 0.87) or interactions (Sex x Condition, *F*(1, 27) = 0.16, *p* = 0.69; Sex x Day, *F*(2, 54) = 0.21, *p* = 0.81, Condition x Day, *F*(2, 54) = 0.38, *p* = 0.68; Sex x Condition x Day, *F*(2, 54) = 0.50, *p* = 0.61) on the preference for the primary imprinting stimulus. The preference for the primary imprinting stimulus was significantly different from chance-level (*t*(30) = -4.24, *p* < 0.001, Cohen’s *d* = 0.76) with an average time spent close to the secondary imprinting stimulus of 63 % (+/- 3.05 SEM).

**Figure 6:**
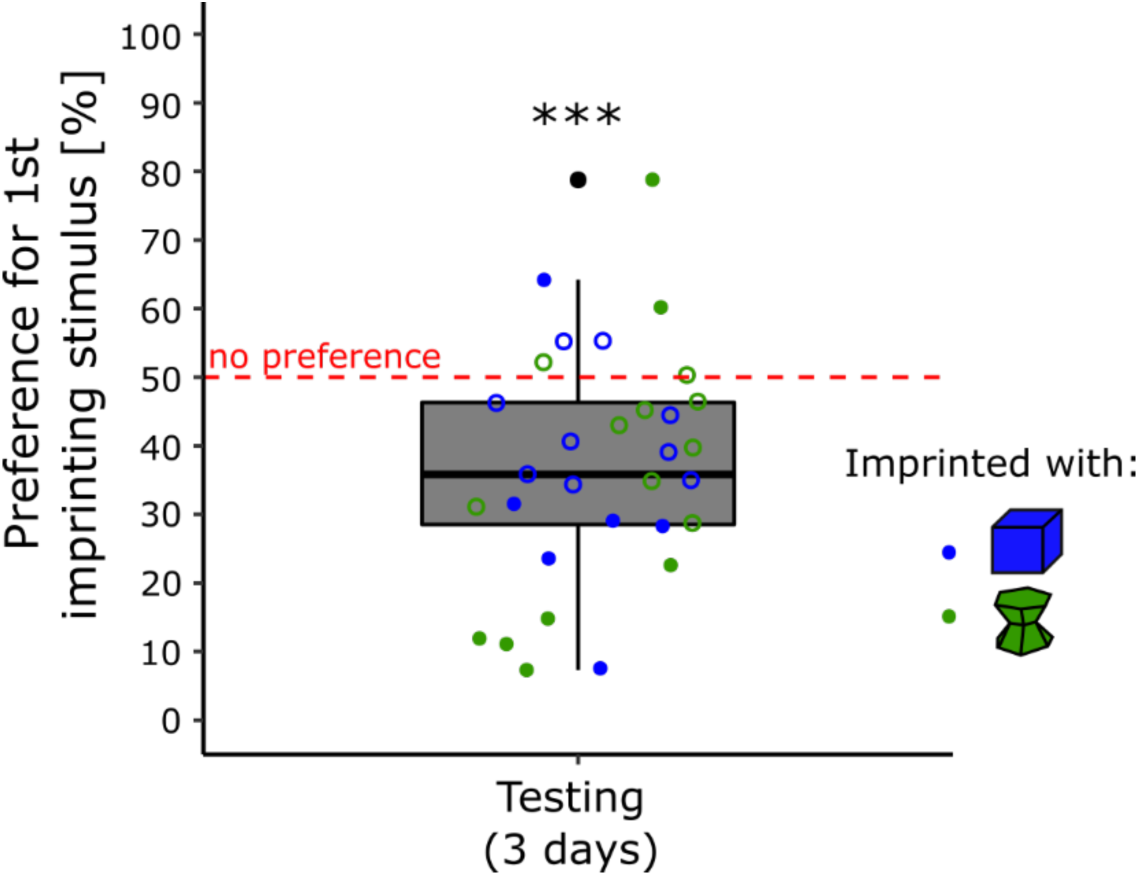
Preference for the primary imprinting stimulus during the testing phase (p < 0.001, ***). The blue dots represent the preference score of the chicks imprinted with the blue stimulus. The green dots represent the preference score of the chicks imprinted with the green stimulus. Filled dots show the individuals having a significant preference while empty dots show the individuals having no preference.

Individual preferences were calculated. In the blue condition, 1 chick (7%) had a significant preference for the imprinting stimulus, 9 (60%) had no preference, and 5 (33%) significantly preferred the unfamiliar stimulus. In the green condition, 2 chicks (13%) had a significant preference for the imprinting stimulus, 9 (56%) had no preference, and 5 (31%) had a significant preference for the unfamiliar stimulus (Table 4 in the supplementary material). Levene’s test showed that the variances of the two conditions were similar (*F*(1, 29) = 2.15, *p* = 0.15).

## Discussion

Due to the difficulties in assessing animals behaviours over prolonged durations, the temporal stability and individual variability of social attachment in filial imprinting have remained unexplored. To understand more about it, we used an automated behavioural tracking and followed the animals’ preferences for familiar and novel stimuli for 6 consecutive days. The temporal stability of the imprinting preferences was investigated by manipulating the duration of the imprinting and the stimuli used. When imprinted for 14 hours over 1 day (Experiment 1), the chicks exhibited a less stable preference for their imprinting stimulus in comparison to when exposed for 42 hours over 3 days (Experiment 2). In fact, after 1 day of imprinting, the filial preferences were disparate between conditions. While the chicks of the blue condition always had a preference for their imprinting stimulus at testing, the chicks of the green condition lost their significant preference for the imprinting stimulus on the fourth testing day. They started to explore more the unfamiliar stimulus (blue stimulus). Since we know that chicks mainly rely on colour to recognize their artificial imprinting objects (Maekawa et al., 2006), this difference confirms previous reports of an advantage of blue over green imprinting stimuli (Kovach, 1971; Salzen et al., 1971; Schaefer & Hess, 2010). In contrast, after 3 days of imprinting, chicks of both conditions had a robust and stable preference for their imprinting objects. Moreover, we excluded the possibility that the difference observed was affected by the time spent close to the imprinting stimuli by showing that bot conditions spent the same amount of time close to their respective stimulus during the imprinting phase.

The preference observed in Experiment 1 for the imprinting stimulus across days in the blue condition and on the first testing day of the green condition indicated that chicks imprinted on their respective stimuli. Nevertheless, 14 hours of imprinting is not sufficient to produce a stable imprinting preference for artificial stimuli. The unlearned preferences are influencing the animals’ filial preferences. Therefore, the decrease of preference for the imprinting stimulus in the green condition could be explained by the fact that the blue stimulus is more attractive to the chicks. This would explain why the animals spend more time close to it to, rather than by a general lack of memory (blue-imprinted chicks steadily remembered and prefered the imprinting stimulus). The difference between blue and green-imprinted chicks is apparent also looking at the individual performances. 63% of the chicks had a preference for the imprinting stimulus, and only 6% had a preference for the novel stimulus in the blue-imprinted chicks. In contrast, only 44% had a preference for the imprinting stimulus, and 19% had a preference for the novel stimulus in the green-imprinted chicks.

Several biochemicals changes associated with imprinting have been described later than 15 hours after the start of the imprinting process, confirming the idea that imprinting might not be fully consolidated in the first day of exposure (Solomonia & McCabe, 2015). Furthermore, the mechanisms responsible for the spontaneous preferences observed in chicks strongly influences the formation of the imprinting memory (Miura & Matsushima, 2016; Miura et al., 2020). In Experiment 1, it seems that after 14 hours of exposure to a stimulus, the imprinting memories are available but not fully consolidated yet. The preferences also seem more plastic after imprinting with less predisposed stimuli.

Hence, because same experience produces different learning outcomes, it appears that predispositions affect both learning and the between-subjects variability in learning, with faster and stronger learning and less variability when subjects are exposed to predisposed stimuli.

The analysis of individual behaviours revealed that some chicks had consistent preferences for unfamiliar stimuli not only at the very beginning of imprinting, as it was instead hypothesised by the Bateson’s model (Bateson, 1973). By increasing the exposure of the chicks to their imprinting objects to 42 hours over 3 days, we observed more robust and stable filial preferences with time for both stimuli (Experiment 2) but still a higher inter-individual variability within the green-imprinted chicks. These results are in line with previous experiments in which preferences for unfamiliar objects have been observed even after 3 days of imprinting (Versace et al., 2006; Versace et al., 2017).

More prolonged imprinting exposure has been associated with stronger preference scores for the imprinting stimulus (Bateson & Jaeckel, 1976; Hess, 1959). Furthermore, our study suggests that the imprinting duration strongly influences the stability of filial preference. After 42 hours over 3 days of exposure to an object, the imprinting memory appears to be consolidated for both artificial stimuli (green and blue). Nonetheless, animals’ predisposed/unlearned preferences for specific stimuli are still, to a lower degree, influencing chicks’ filial preferences. The variability within the green condition (less predisposed colour) was three-time higher than in the blue condition. While almost all chicks (87.5%) showed a strong preference for their imprinting objects in the blue condition, 37.5% did not prefer their imprinting stimulus in the green condition (12.5% prefered the unfamiliar stimulus and 25% had no preference).

The evidence that prolonged exposure to an object leads to more stable preferences in time is convincing and in line with previous evidence (Bateson & Jaeckel, 1976; Bolhuis et al., 2000). Nevertheless, the ontogenetic stage at which the preferences were tested could have influenced filial preferences. In the third experiment, we assessed whether this was the case. As in the first experiment, both conditions (blue and green) were exposed to their respective objects for 14 hours (day 1), but this time, their filial preference was tested from day 4 to day 6, after exposure to a novel object on day 2 and 3 (this prevented a complete ‘social’ deprivation). Similarly to what observed in the first experiment (short imprinting duration), the filial preferences observed were different between conditions. In the blue condition, all the individuals preferred their imprinting object, showing that the memory of the imprinting stimulus lasted although chicks had been detached by the initial stimulus for days. At the same time, preferences among individuals of the green conditions were disparate with 13% of the individuals preferring the imprinting object, 37% showing no preferences and even 50% showing a preference for the novel object. Interestingly, the preferences observed here were not wholly similar to the first experiment. The preferences observed in both conditions were stable in time. Then, one could argue that the filial preferences observed were the result of a lack of memory, but the different patterns of preference between conditions and the literature suggest otherwise. In the case of a memory loss, chicks would have either approached the more attractive stimulus (blue object) or not chosen any. However, the results showed both patterns depending on the primary imprinting stimulus used. Moreover, studies exploring successive imprinting always described a recall of the primary imprinting object (Bolhuis & Bateson, 1990; Salzen & Meyer, 1968).

In Experiment 4, we assessed whether chicks had a preference for their primary imprinting stimulus in comparison to their secondary imprinting stimulus during the testing phase. Both conditions showed a similar preference for the secondary imprinting stimulus. As previously shown, chicks can imprint on multiple objects (Boakes & Panter, 1985; Bolhuis & Trooster, 1988). Furthermore, a preference for a primary imprinting stimulus can be reversed after prolonged exposure with a secondary imprinting object (Cherfas & Scott, 1981), which is line with the experimental settings used here (one day of primary imprinting and two days of secondary imprinting). It is then very likely that the filial bond formed with the secondary imprinting object has influenced the filial preferences of the chicks toward their primary imprinting stimulus.

In all experiments, the filial imprinting preferences were all pointing in the same direction: Overall, chicks of the blue condition (where blue is a more predisposed colour) had a more robust and stable preference in time for their imprinting stimulus than the chicks of the green condition (where green is a less predisposed colour). The differences between conditions were not the result of the time spent close their respective objects during imprinting, given that chicks engaged with the imprinting stimuli for the same amount of time. This strongly suggests that some features of the objects (e.g. colour) are more efficient for the formation of filial imprinting preferences.

Altogether, our results indicate that the temporal stability of filial imprinting preferences is influenced by the amount of experience (exposure duration and successive imprinting) and spontaneous preferences (predispositions). Moreover, using automated tracking for assessing chicks’ behaviour for several days, we show that chicks with similar experiences can have steady and robust idiosyncratic differences in their preferences for familiar vs novel stimuli. Some chicks consistently preferred to approach their imprinting stimulus while others preferred the unfamiliar stimulus, even if they had the same experience. Moreover, this consistent inter-individual variability (a phenomenon already documented in other model systems, Buchanan, Kain, & de Bivort, 2015; Honegger, Smith, Churgin, Turner, & de Bivort, 2019; Kain, Stokes, & de Bivort, 2012; Kain et al., 2015; Linneweber et al., 2020; Versace et al., 2020) was modulated by the animals’ unlearned preferences. Further studies should clarify whether these differences stem from genetic variability or derive from stochasticity in the course of development (Mitchell, 2018), as well as their neurobiological basis.

## Data accessibility

The datasets (.csv) are available on Fig **Share** (10.6084/m9.figshare.12074565).

## Authors’ contribution

B.S.L. conceived and performed the research, collected and analysed the data and drafted the manuscript. D.R. and M.J. helped to collect and analyse the data. G.V. and E.V. conceived the experiment and helped in drafting the manuscript. E.V. supervised the project. All authors gave final approval for publication and are accountable for the work performed.

