## Supplemental Tables for "Stability and individual variability of social attachment in imprinting"

Table 1: Table showing the results of the t-tests on the preference for the imprinting stimulus against chance-level (50%) for each individual of the first experiment.

| | **Imprinting** | | | | | | | **Testing** | | | | | |
|---|---|---|---|---|---|---|---|---|---|---|---|---|---|
| | | | | | 95% CI | | | | | | 95% CI | | |
| | Ind. | *df* | t | *Mean* | *Lower* | *Upper* | *Significance* | *df* | t | *Mean* | *Lower* | *Upper* | *Significance* |
| Blue condition | 1 | 6 | 10.56306 | 93.90871 | 83.73734 | 104.0801 | < 0.001, *** | 119 | 7.149224 | 69.1932 | 63.87732 | 74.50909 | < 0.001, *** |
| | 2 | 6 | 12.55208 | 93.80412 | 85.26492 | 102.3433 | < 0.001, *** | 119 | 10.2481 | 73.77989 | 69.18522 | 78.37455 | < 0.001, *** |
| | 3 | 6 | 72.84424 | 97.68572 | 96.0839 | 99.28753 | < 0.001, *** | 119 | 0.949654 | 52.9253 | 46.82583 | 59.02477 | 0.34 |
| | 4 | 6 | 10.59125 | 92.89619 | 82.98582 | 102.8066 | < 0.001, *** | 119 | 17.37157 | 85.19431 | 81.18268 | 89.20593 | < 0.001, *** |
| | 5 | 6 | 25.9876 | 95.5669 | 91.27646 | 99.85734 | < 0.001, *** | 119 | 7.144761 | 65.40466 | 61.13541 | 69.67392 | < 0.001, *** |
| | 6 | 6 | 36.21517 | 97.718 | 94.49389 | 100.9421 | < 0.001, *** | 119 | -0.71778 | 47.3247 | 39.94447 | 54.70494 | 0.47 |
| | 7 | 6 | 42.27593 | 97.69005 | 94.92977 | 100.4503 | < 0.001, *** | 119 | 0.207854 | 50.56639 | 45.17076 | 55.96202 | 0.84 |
| | 8 | 6 | 10.22111 | 94.37024 | 83.7481 | 104.9924 | < 0.001, *** | 119 | -0.00392 | 49.99007 | 44.97838 | 55.00177 | 1.00 |
| | 9 | 6 | 3.203028 | 86.47754 | 58.611 | 114.3441 | < 0.05, * | 119 | 3.718203 | 61.1604 | 55.21701 | 67.10378 | < 0.001, *** |
| | 10 | 6 | 44.05424 | 98.63238 | 95.93119 | 101.3336 | < 0.001, *** | 119 | 1.488381 | 55.72576 | 48.10837 | 63.34315 | 0.14 |
| | 11 | 6 | 115.4368 | 98.99096 | 97.9525 | 100.0294 | < 0.001, *** | 119 | 13.86966 | 77.71457 | 73.7579 | 81.67123 | < 0.001, *** |
| | 12 | 6 | 6.139136 | 91.58199 | 75.00841 | 108.1556 | < 0.001, *** | 119 | 3.008457 | 59.40851 | 53.21604 | 65.60099 | < 0.01, ** |
| | 13 | 6 | 17.96936 | 96.63756 | 90.28686 | 102.9883 | < 0.001, *** | 119 | -2.1389 | 43.40938 | 37.30807 | 49.5107 | < 0.05, * |
| | 14 | 6 | 36.07158 | 98.24851 | 94.97557 | 101.5214 | < 0.001, *** | 119 | 2.746845 | 57.78278 | 52.17246 | 63.3931 | < 0.01, ** |
| | 15 | 6 | 7.515497 | 88.52483 | 75.98183 | 101.0678 | < 0.001, *** | 117 | 4.308517 | 63.01653 | 57.03337 | 68.9997 | < 0.001, *** |
| | 16 | 6 | 23.48606 | 94.97415 | 90.28849 | 99.65982 | < 0.001, *** | 119 | 23.39976 | 85.09009 | 82.12074 | 88.05943 | < 0.001, *** |
| Green condition | 17 | 6 | 8.82454 | 94.52126 | 82.17619 | 106.8663 | < 0.001, *** | 119 | 1.73518 | 55.40616 | 49.23692 | 61.5754 | 0.09 |
| | 18 | 6 | 32.41124 | 97.48144 | 93.89679 | 101.0661 | < 0.001, *** | 119 | 2.88149 | 59.2249 | 52.88574 | 65.56406 | < 0.01, ** |
| | 19 | 6 | 314.0042 | 99.62547 | 99.23876 | 100.0122 | < 0.001, *** | 119 | 1.080542 | 53.79864 | 46.83761 | 60.75966 | 0.28 |
| | 20 | 6 | 16.22367 | 93.34748 | 86.80966 | 99.8853 | < 0.001, *** | 119 | -1.56128 | 44.61696 | 37.78988 | 51.44404 | 0.12 |
| | 21 | 6 | 19.87124 | 96.52046 | 90.79201 | 102.2489 | < 0.001, *** | 119 | -3.08236 | 39.26922 | 32.3758 | 46.16263 | < 0.01, ** |
| | 22 | 6 | 42.58746 | 97.13972 | 94.43126 | 99.84819 | < 0.001, *** | 119 | 1.176899 | 53.51557 | 47.60072 | 59.43043 | 0.24 |
| | 23 | 6 | 117.8321 | 99.2683 | 98.24519 | 100.2914 | < 0.001, *** | 119 | -5.40546 | 33.92363 | 28.03461 | 39.81264 | < 0.001, *** |
| | 24 | 6 | 151.2527 | 99.32711 | 98.52912 | 100.1251 | < 0.001, *** | 119 | 2.956339 | 60.64189 | 53.51415 | 67.76962 | < 0.01, ** |
| | 25 | 6 | 25.18918 | 96.89309 | 92.33783 | 101.4483 | < 0.001, *** | 119 | 0.638648 | 52.11696 | 45.55342 | 58.68049 | 0.52 |
| | 26 | 6 | 238.2557 | 99.4259 | 98.91829 | 99.93351 | < 0.001, *** | 119 | 0.764078 | 52.56237 | 45.92201 | 59.20274 | 0.45 |
| | 27 | 6 | 113.5141 | 98.41019 | 97.36666 | 99.45372 | < 0.001, *** | 119 | 45.34628 | 93.31335 | 91.42202 | 95.20468 | < 0.001, *** |
| | 28 | 6 | 47.93681 | 98.06098 | 95.60773 | 100.5142 | < 0.001, *** | 119 | -2.54728 | 41.56173 | 35.00232 | 48.12113 | < 0.05, * |
| | 29 | 6 | 42.65225 | 97.76618 | 95.02589 | 100.5065 | < 0.001, *** | 119 | 3.43273 | 60.13391 | 54.28837 | 65.97944 | < 0.001, *** |
| | 30 | 6 | 132.7135 | 98.95841 | 98.05574 | 99.86108 | < 0.001, *** | 119 | 4.825088 | 63.07635 | 57.71013 | 68.44257 | < 0.001, *** |
| | 31 | 6 | 64.58975 | 98.33763 | 96.50641 | 100.1689 | < 0.001, *** | 119 | 7.15215 | 71.70061 | 65.69271 | 77.70851 | < 0.001, *** |
| | 32 | 6 | 49.00067 | 97.28253 | 94.92141 | 99.64364 | < 0.001, *** | 119 | 13.82374 | 80.72518 | 76.32414 | 85.12623 | < 0.001, *** |

Table 2: Table showing the results of the t-tests on the preference for the imprinting stimulus against chance-level (50%) for each individual of the second experiment.

**Experiment 2**

| | **Imprinting** | | | | | | | **Testing** | | | | | |
|---|---|---|---|---|---|---|---|---|---|---|---|---|---|
| | | | | | 95% CI | | | | | | 95% CI | | |
| | Ind. | *df* | t | *Mean* | *Lower* | *Upper* | *Significance* | *df* | t | *Mean* | *Lower* | *Upper* | *Significance* |
| Blue condition | 1 | 32 | 16.57942 | 91.3977 | 86.31162 | 96.48379 | < 0.001, *** | 71 | 5.892126 | 72.46538 | 64.86291 | 80.06784 | < 0.001, *** |
| | 2 | 32 | 9.879792 | 89.03787 | 80.98937 | 97.08638 | < 0.001, *** | 68 | 8.347324 | 75.7069 | 69.56154 | 81.85226 | < 0.001, *** |
| | 3 | 32 | 5.571881 | 79.5776 | 68.76481 | 90.3904 | < 0.001, *** | 71 | 4.590961 | 66.34672 | 59.24702 | 73.44642 | < 0.001, *** |
| | 4 | 32 | 39.39346 | 94.46552 | 92.16632 | 96.76472 | < 0.001, *** | 68 | 6.719652 | 75.18352 | 67.70502 | 82.66202 | < 0.001, *** |
| | 5 | 32 | 135.3361 | 98.60293 | 97.87141 | 99.33445 | < 0.001, *** | 71 | 11.09174 | 79.36821 | 74.08874 | 84.64769 | < 0.001, *** |
| | 6 | 32 | 53.99766 | 96.88197 | 95.11346 | 98.65048 | < 0.001, *** | 71 | 30.64861 | 95.75983 | 92.78277 | 98.73688 | < 0.001, *** |
| | 7 | 32 | 14.9338 | 94.46806 | 88.40273 | 100.5334 | < 0.001, *** | 71 | 5.979519 | 72.9727 | 65.31217 | 80.63322 | < 0.001, *** |
| | 8 | 32 | 30.60358 | 95.32222 | 92.30563 | 98.3388 | < 0.001, *** | 71 | 1.326022 | 55.37028 | 47.29497 | 63.4456 | 0.19 |
| | 9 | 32 | 29.97684 | 95.02616 | 91.96663 | 98.0857 | < 0.001, *** | 71 | 6.598297 | 72.69741 | 65.83846 | 79.55635 | < 0.001, *** |
| | 10 | 32 | 29.56029 | 94.16985 | 91.1262 | 97.21349 | < 0.001, *** | 71 | 3.016844 | 63.3243 | 54.51778 | 72.13081 | < 0.01, ** |
| | 11 | 32 | 48.04413 | 97.75639 | 95.73165 | 99.78112 | < 0.001, *** | 71 | 8.298398 | 76.13616 | 69.85615 | 82.41617 | < 0.001, *** |
| | 12 | 32 | 44.87811 | 96.49052 | 94.3804 | 98.60064 | < 0.001, *** | 71 | 18.9641 | 90.20689 | 85.97942 | 94.43437 | < 0.001, *** |
| | 13 | 32 | 6.555537 | 83.07705 | 72.79936 | 93.35473 | < 0.001, *** | 71 | 4.072488 | 68.20719 | 59.29271 | 77.12167 | < 0.001, *** |
| | 14 | 32 | 10.78176 | 85.98622 | 79.18756 | 92.78488 | < 0.001, *** | 71 | 10.40145 | 77.55147 | 72.26989 | 82.83304 | < 0.001, *** |
| | 15 | 32 | 38.62432 | 94.57548 | 92.2247 | 96.92626 | < 0.001, *** | 71 | 0.611928 | 52.41612 | 44.54328 | 60.28897 | 0.54 |
| | 16 | 32 | 22.89494 | 93.97139 | 90.05931 | 97.88347 | < 0.001, *** | 71 | 15.34234 | 85.80319 | 81.15008 | 90.45629 | < 0.001, *** |
| Green condition | 17 | 32 | 45.88517 | 97.60693 | 95.49356 | 99.72029 | < 0.001, *** | 71 | 0.619212 | 52.68119 | 44.04742 | 61.31495 | 0.54 |
| | 18 | 32 | 54.03456 | 97.47238 | 95.68282 | 99.26194 | < 0.001, *** | 71 | 2.289018 | 58.08895 | 51.04274 | 65.13517 | < 0.05, * |
| | 19 | 32 | 7.263146 | 85.20741 | 75.33357 | 95.08125 | < 0.001, *** | 71 | 4.851796 | 68.50595 | 60.90056 | 76.11134 | < 0.001, *** |
| | 20 | 32 | 17.49157 | 92.124 | 87.21856 | 97.02943 | < 0.001, *** | 71 | -0.01021 | 49.96083 | 42.30772 | 57.61394 | 0.99 |
| | 21 | 32 | 82.41629 | 98.13098 | 96.94142 | 99.32055 | < 0.001, *** | 71 | 20.25957 | 89.09465 | 85.24696 | 92.94234 | < 0.001, *** |
| | 22 | 32 | 5.824866 | 83.65738 | 71.88752 | 95.42723 | < 0.001, *** | 71 | 1.932702 | 57.36142 | 49.76674 | 64.95609 | 0.06 |
| | 23 | 32 | 37.36719 | 96.79376 | 94.24297 | 99.34455 | < 0.001, *** | 71 | 10.31046 | 78.2492 | 72.78607 | 83.71232 | < 0.001, *** |
| | 24 | 32 | 23.8663 | 93.2849 | 89.59063 | 96.97916 | < 0.001, *** | 71 | 7.809482 | 74.66952 | 68.37082 | 80.96823 | < 0.001, *** |
| | 25 | 32 | 32.89396 | 95.74547 | 92.91272 | 98.57823 | < 0.001, *** | 71 | -8.05909 | 24.78685 | 18.54872 | 31.02497 | < 0.001, *** |
| | 26 | 32 | 82.48646 | 97.35087 | 96.18158 | 98.52016 | < 0.001, *** | 71 | 6.998888 | 70.9132 | 64.95515 | 76.87125 | < 0.001, *** |
| | 27 | 32 | 17.95306 | 92.27538 | 87.47887 | 97.0719 | < 0.001, *** | 71 | -2.05476 | 43.54265 | 37.27644 | 49.80886 | < 0.05, * |
| | 28 | 32 | 47.52607 | 95.27593 | 93.33544 | 97.21643 | < 0.001, *** | 71 | 2.229398 | 56.69627 | 50.70722 | 62.68532 | < 0.05, * |
| | 29 | 32 | 22.94583 | 94.34859 | 90.4117 | 98.28547 | < 0.001, *** | 71 | -1.17044 | 43.55872 | 32.58546 | 54.53199 | 0.25 |
| | 30 | 32 | 148.6678 | 98.57376 | 97.90824 | 99.23928 | < 0.001, *** | 71 | 86.81891 | 98.278 | 97.16922 | 99.38679 | < 0.001, *** |
| | 31 | 32 | 26.5803 | 96.59359 | 93.02297 | 100.1642 | < 0.001, *** | 71 | 9.034338 | 80.66798 | 73.89933 | 87.43662 | < 0.001, *** |
| | 32 | 31 | 24.94614 | 94.1863 | 90.57377 | 97.79882 | < 0.001, *** | 71 | 15.2027 | 84.42542 | 79.91028 | 88.94056 | < 0.001, *** |

Table 3: Table showing the results of the t-tests on the preference for the primary imprinting stimulus against chance-level (50%) for each individual of the third experiment.

**Experiment 3**

| | | Primary Imprinting | | | | | | Secondary Imprinting | | | | | | Testing | | | | | |
|---|---|---|---|---|---|---|---|---|---|---|---|---|---|---|---|---|---|---|---|
| | | | | | 95% CI | | | | | | 95% CI | | | | | | 95% CI | | |
| | Ind. | df | t | Mean | Lower | Upper | Significance | df | t | Mean | Lower | Upper | Significance | df | t | Mean | Lower | Upper | Significance |
| Blue condition | 1 | 6 | 34.8885 | 98.03451 | 94.6656 | 101.4034 | < 0.001, *** | 25 | 2.792299 | 67.81053 | 54.67387 | 80.94719 | < 0.01, ** | 69 | 6.734117 | 71.31123 | 64.99789 | 77.62456 | < 0.001, *** |
| | 2 | 6 | 11.35221 | 95.00117 | 85.30139 | 104.701 | < 0.001, *** | 25 | 110.6497 | 99.08529 | 98.17165 | 99.99892 | < 0.001, *** | 71 | 13.34891 | 84.89516 | 79.68283 | 90.10749 | < 0.001, *** |
| | 3 | 6 | 24.7871 | 96.90038 | 92.27051 | 101.5303 | < 0.001, *** | 25 | 12.02614 | 92.0413 | 84.84151 | 99.24109 | < 0.001, *** | 71 | 11.95003 | 84.04508 | 78.36443 | 89.72573 | < 0.001, *** |
| | 4 | 6 | 36.14095 | 97.56757 | 94.34702 | 100.7881 | < 0.001, *** | 25 | 240.8659 | 99.67001 | 99.2453 | 100.0947 | < 0.001, *** | 71 | 112.7937 | 98.3068 | 97.45285 | 99.16076 | < 0.001, *** |
| | 5 | 6 | 144.7477 | 99.32547 | 98.49164 | 100.1593 | < 0.001, *** | 25 | 170.8064 | 98.99561 | 98.40484 | 99.58639 | < 0.001, *** | 71 | 14.26569 | 87.59051 | 82.33641 | 92.84461 | < 0.001, *** |
| | 6 | 6 | 368.1679 | 99.55114 | 99.22182 | 99.88047 | < 0.001, *** | 25 | 13.30654 | 90.7989 | 84.48419 | 97.1136 | < 0.001, *** | 71 | 6.898919 | 72.53382 | 66.02104 | 79.0466 | < 0.001, *** |
| | 7 | 6 | 94.04666 | 98.5095 | 97.24738 | 99.77162 | < 0.001, *** | 25 | 240.1073 | 99.24072 | 98.81835 | 99.66308 | < 0.001, *** | 71 | 15.96917 | 84.1513 | 79.8871 | 88.4155 | < 0.001, *** |
| | 8 | 6 | 25.30403 | 96.35184 | 91.86959 | 100.8341 | < 0.001, *** | 25 | 17.72484 | 95.14894 | 89.90286 | 100.395 | < 0.001, *** | 71 | 31.01566 | 96.11332 | 93.14878 | 99.07787 | < 0.001, *** |
| | 9 | 6 | 41.99048 | 97.90402 | 95.1125 | 100.6955 | < 0.001, *** | 25 | 21.92715 | 95.55536 | 91.27651 | 99.83421 | < 0.001, *** | 71 | 9.225992 | 78.43108 | 72.28649 | 84.57568 | < 0.001, *** |
| | 10 | 6 | 71.53717 | 98.8824 | 97.21039 | 100.5544 | < 0.001, *** | 25 | 5.25937 | 76.67114 | 66.22688 | 87.1154 | < 0.001, *** | 71 | 7.619617 | 74.27038 | 67.91917 | 80.62159 | < 0.001, *** |
| | 11 | 6 | 27.75394 | 97.35179 | 93.17705 | 101.5265 | < 0.001, *** | 25 | 24.78711 | 93.51608 | 89.90036 | 97.13179 | < 0.001, *** | 69 | 10.21703 | 80.23803 | 74.33385 | 86.14221 | < 0.001, *** |
| | 12 | 6 | 17.04717 | 97.12782 | 90.3632 | 103.8924 | < 0.001, *** | 24 | 37.76702 | 97.402 | 94.81157 | 99.99243 | < 0.001, *** | 71 | 65.06488 | 96.97838 | 95.53871 | 98.41806 | < 0.001, *** |
| | 13 | 6 | 148.3466 | 99.05155 | 98.24247 | 99.86063 | < 0.001, *** | 24 | 88.0002 | 97.13924 | 96.03367 | 98.24481 | < 0.001, *** | 71 | 5.867247 | 68.20542 | 62.01844 | 74.39241 | < 0.001, *** |
| | 14 | 6 | 76.37756 | 98.61525 | 97.05776 | 100.1727 | < 0.001, *** | 24 | 51.41571 | 98.49681 | 96.55008 | 100.4435 | < 0.001, *** | 71 | 32.6507 | 93.84947 | 91.17163 | 96.52731 | < 0.001, *** |
| | 15 | 6 | 38.65896 | 98.33786 | 95.27832 | 101.3974 | < 0.001, *** | 25 | 17.29401 | 93.74931 | 88.53922 | 98.95941 | < 0.001, *** | 71 | 3.701949 | 65.14853 | 56.98923 | 73.30783 | < 0.001, *** |
| | 16 | 6 | 78.00533 | 97.85833 | 96.35708 | 99.35957 | < 0.001, *** | 25 | 34.89179 | 96.07216 | 93.35269 | 98.79164 | < 0.001, *** | 71 | 29.9753 | 96.0485 | 92.98537 | 99.11162 | < 0.001, *** |
| | 17 | 6 | 73.79672 | 96.83951 | 95.28643 | 98.39259 | < 0.001, *** | 25 | 93.45334 | 97.51313 | 96.46603 | 98.56024 | < 0.001, *** | 71 | 5.528457 | 72.13149 | 64.14935 | 80.11364 | < 0.001, *** |
| Green condition | 18 | 6 | 5.355679 | 86.70097 | 69.93296 | 103.469 | < 0.01, ** | 25 | 11.5473 | 91.85069 | 84.38634 | 99.31503 | < 0.001, *** | 71 | -4.59227 | 29.27618 | 20.27799 | 38.27436 | < 0.001, *** |
| | 19 | 6 | 46.96131 | 98.22628 | 95.71346 | 100.7391 | < 0.001, *** | 25 | 185.4095 | 98.86848 | 98.32565 | 99.41131 | < 0.001, *** | 71 | 0.627835 | 52.60857 | 44.324 | 60.89314 | 0.53 |
| | 20 | 6 | 42.79792 | 97.47745 | 94.76299 | 100.1919 | < 0.001, *** | 25 | 14.49014 | 93.27228 | 87.12183 | 99.42273 | < 0.001, *** | 71 | -3.31322 | 35.16257 | 26.23321 | 44.09194 | < 0.01, ** |
| | 21 | 6 | 15.06915 | 95.84237 | 88.39853 | 103.2862 | < 0.001, *** | 25 | 61.66778 | 96.14787 | 94.60666 | 97.68909 | < 0.001, *** | 71 | 0.954228 | 54.60245 | 44.98523 | 64.21968 | 0.34 |
| | 22 | 6 | 11.8785 | 91.0898 | 82.6255 | 99.55409 | < 0.001, *** | 25 | 11.31651 | 92.08613 | 84.4267 | 99.74555 | < 0.001, *** | 71 | -1.13755 | 45.36763 | 37.24779 | 53.48746 | 0.26 |
| | 23 | 6 | 307.1818 | 99.59291 | 99.19787 | 99.98795 | < 0.001, *** | 25 | 35.43525 | 96.01402 | 93.33963 | 98.68841 | < 0.001, *** | 71 | -4.0498 | 34.3549 | 26.65193 | 42.05787 | < 0.001, *** |
| | 24 | 6 | 116.9607 | 98.70982 | 97.69077 | 99.72886 | < 0.001, *** | 25 | 32.78368 | 97.10675 | 94.1474 | 100.0661 | < 0.001, *** | 71 | 7.417937 | 78.60812 | 70.91825 | 86.29799 | < 0.001, *** |
| | 25 | 6 | 120.426 | 99.16922 | 98.17016 | 100.1683 | < 0.001, *** | 25 | 28.82765 | 93.57097 | 90.45812 | 96.68381 | < 0.001, *** | 71 | 3.214488 | 62.26888 | 54.65851 | 69.87925 | < 0.01, ** |
| | 26 | 6 | 9.753007 | 93.85507 | 82.85236 | 104.8578 | < 0.001, *** | 25 | 15.52545 | 93.03565 | 87.32672 | 98.74457 | < 0.001, *** | 71 | -11.8335 | 15.37612 | 9.541998 | 21.21024 | < 0.001, *** |
| | 27 | 6 | 11.26724 | 95.424 | 85.55924 | 105.2887 | < 0.001, *** | 25 | 37.33564 | 94.4846 | 92.03071 | 96.9385 | < 0.001, *** | 71 | -2.82933 | 36.10333 | 26.30978 | 45.89687 | < 0.01, ** |
| | 28 | 6 | 4.219447 | 88.27241 | 66.07774 | 110.4671 | < 0.01, ** | 25 | 25.63091 | 95.08547 | 91.46269 | 98.70826 | < 0.001, *** | 71 | 0.421784 | 51.55349 | 44.20952 | 58.89747 | 0.67 |
| | 29 | 6 | 190.2468 | 99.51347 | 98.87663 | 100.1503 | < 0.001, *** | 25 | 4.73435 | 78.2326 | 65.95084 | 90.51435 | < 0.001, *** | 71 | -0.19856 | 49.13937 | 40.49694 | 57.7818 | 0.84 |
| | 30 | 6 | 81.48075 | 98.89262 | 97.42435 | 100.3609 | < 0.001, *** | 25 | 31.06984 | 95.75915 | 92.7259 | 98.79241 | < 0.001, *** | 71 | -1.16071 | 44.37984 | 34.72514 | 54.03454 | 0.25 |
| | 31 | 6 | 132.4911 | 99.23591 | 98.3266 | 100.1452 | < 0.001, *** | 25 | 31.72703 | 95.4731 | 92.52125 | 98.42496 | < 0.001, *** | 71 | -6.18444 | 25.65792 | 17.80971 | 33.50613 | < 0.001, *** |
| | 32 | 6 | 12.32503 | 95.28547 | 86.29486 | 104.2761 | < 0.001, *** | 25 | 39.79303 | 97.5003 | 95.04186 | 99.95874 | < 0.001, *** | 71 | -3.6172 | 34.81925 | 26.45103 | 43.18747 | < 0.001, *** |
| | 33 | 6 | 718.2703 | 99.82502 | 99.65528 | 99.99476 | < 0.001, *** | 25 | 5.226489 | 81.11336 | 68.8529 | 93.37382 | < 0.001, *** | 71 | -4.52271 | 29.90533 | 21.04611 | 38.76454 | < 0.001, *** |

Table 4: Table showing the results of the t-tests on the preference for the primary imprinting stimulus against chance-level (50%) for each individual of the fourth experiment.

**Experiment 4**

| | | Primary Imprinting | | | | | | Secondary Imprinting | | | | | | Testing | | | | | |
|---|---|---|---|---|---|---|---|---|---|---|---|---|---|---|---|---|---|---|---|
| | | | | | 95% CI | | | | | | 95% CI | | | | | | 95% CI | | |
| | Ind. | df | t | Mean | Lower | Upper | Significance | df | t | Mean | Lower | Upper | Significance | df | t | Mean | Lower | Upper | Significance |
| Blue condition | 1 | 6 | 1.667083 | 74.10815 | 38.72267 | 109.4936 | 0.15 | 25 | 96.44095 | 98.79959 | 97.75745 | 99.84172 | < 0.001, *** | 71 | -6.94404 | 23.5949 | 16.01282 | 31.17698 | < 0.001, *** |
| | 2 | 6 | 11.59723 | 95.68107 | 86.04277 | 105.3194 | < 0.001, *** | 25 | 51.25137 | 97.31369 | 95.41238 | 99.21499 | < 0.001, *** | 71 | -1.37876 | 44.48719 | 36.51462 | 52.45976 | 0.17 |
| | 3 | 6 | 59.25317 | 98.36152 | 96.36438 | 100.3586 | < 0.001, *** | 25 | 25.92225 | 96.49728 | 92.80305 | 100.1915 | < 0.001, *** | 71 | -6.32418 | 29.06558 | 22.4652 | 35.66597 | < 0.001, *** |
| | 4 | 6 | 24.08176 | 95.11452 | 90.53051 | 99.69854 | < 0.001, *** | 25 | 23.07775 | 92.34636 | 88.56722 | 96.12549 | < 0.001, *** | 71 | 1.507145 | 55.28551 | 48.29281 | 62.27821 | 0.14 |
| | 5 | 6 | 32.22804 | 96.68663 | 93.14195 | 100.2313 | < 0.001, *** | 25 | 74.50806 | 98.18152 | 96.8497 | 99.51335 | < 0.001, *** | 55 | -3.22713 | 35.83518 | 27.03883 | 44.63153 | < 0.01, ** |
| | 6 | 6 | 7.697672 | 87.57496 | 75.63075 | 99.51917 | < 0.001, *** | 25 | 7.364524 | 86.59695 | 76.36236 | 96.83153 | < 0.001, *** | 71 | -4.04254 | 35.01283 | 27.62056 | 42.4051 | < 0.001, *** |
| | 7 | 6 | 16.57924 | 96.22006 | 89.3985 | 103.0416 | < 0.001, *** | 25 | 7.643681 | 84.40385 | 75.13396 | 93.67373 | < 0.001, *** | 71 | 3.509582 | 64.21798 | 56.14014 | 72.29583 | < 0.001, *** |
| | 8 | 6 | 40.65207 | 97.49352 | 94.63481 | 100.3522 | < 0.001, *** | 25 | 16.65168 | 93.43412 | 88.06204 | 98.80621 | < 0.001, *** | 71 | 1.285933 | 55.20567 | 47.13385 | 63.27749 | 0.20 |
| | 9 | 6 | 44.95973 | 98.50534 | 95.86546 | 101.1452 | < 0.001, *** | 25 | 260.5275 | 99.29463 | 98.90495 | 99.68432 | < 0.001, *** | 71 | -29.8504 | 7.547392 | 4.711652 | 10.38313 | < 0.001, *** |
| | 10 | 6 | 36.69526 | 97.37005 | 94.21132 | 100.5288 | < 0.001, *** | 25 | 57.42403 | 95.85695 | 94.21227 | 97.50163 | < 0.001, *** | 71 | -4.15285 | 39.10878 | 33.87949 | 44.33808 | < 0.001, *** |
| | 11 | 6 | 6.446881 | 90.25624 | 74.97699 | 105.5355 | < 0.001, *** | 25 | 11.64442 | 90.08329 | 82.9938 | 97.17279 | < 0.001, *** | 71 | -6.19819 | 28.31417 | 21.33789 | 35.29045 | < 0.001, *** |
| | 12 | 6 | 2.456265 | 84.73215 | 50.13226 | 119.332 | < 0.05, * | 25 | 78.88674 | 98.10759 | 96.85161 | 99.36356 | < 0.001, *** | 71 | -4.77836 | 31.57167 | 23.88177 | 39.26156 | < 0.001, *** |
| | 13 | 6 | 9.521323 | 90.31368 | 79.95335 | 100.674 | < 0.001, *** | 25 | 19.11081 | 93.99483 | 89.25358 | 98.73607 | < 0.001, *** | 71 | -4.2893 | 34.3567 | 27.08468 | 41.62871 | < 0.001, *** |
| | 14 | 6 | 20.24995 | 95.8163 | 90.28006 | 101.3525 | < 0.001, *** | 25 | 12.76112 | 88.3252 | 82.13983 | 94.51057 | < 0.001, *** | 55 | -0.8326 | 46.24064 | 37.192 | 55.28927 | 0.41 |
| | 15 | 6 | 11.36249 | 93.52392 | 84.15105 | 102.8968 | < 0.001, *** | 25 | 11.09045 | 92.29174 | 84.438 | 100.1455 | < 0.001, *** | 71 | -1.95 | 40.62346 | 31.03563 | 50.2113 | 0.06 |
| | 16 | 6 | 216.3313 | 99.35369 | 98.79545 | 99.91192 | < 0.001, *** | 25 | 105.1032 | 97.8048 | 96.86804 | 98.74155 | < 0.001, *** | removed from the analysis | | | | | |
| | 17 | 6 | 38.88033 | 98.23019 | 95.19485 | 101.2655 | < 0.001, *** | 25 | 28.97368 | 95.67032 | 92.42394 | 98.91671 | < 0.001, *** | | | | | | |
| Green condition | 18 | 6 | 30.62308 | 95.81293 | 92.15228 | 99.47357 | < 0.001, *** | 25 | 2.677698 | 67.9004 | 54.13239 | 81.66841 | < 0.05, * | 67 | 2.486425 | 60.19257 | 52.01036 | 68.37479 | < 0.05, * |
| | 19 | 6 | 62.29047 | 96.96666 | 95.12171 | 98.81162 | < 0.001, *** | 25 | 24.1507 | 91.59655 | 88.04925 | 95.14384 | < 0.001, *** | 67 | 10.15782 | 78.77264 | 73.11883 | 84.42646 | < 0.001, *** |
| | 20 | 6 | 68.25934 | 98.63737 | 96.89385 | 100.3809 | < 0.001, *** | 25 | 24.04967 | 92.22926 | 88.61288 | 95.84565 | < 0.001, *** | 71 | -5.55651 | 28.73861 | 21.10899 | 36.36822 | < 0.001, *** |
| | 21 | 6 | 131.9186 | 99.41859 | 98.50195 | 100.3352 | < 0.001, *** | 25 | 91.6703 | 97.41196 | 96.34677 | 98.47716 | < 0.001, *** | 71 | -5.0149 | 31.08796 | 23.56846 | 38.60745 | < 0.001, *** |
| | 22 | 6 | 29.11512 | 96.34784 | 92.45264 | 100.243 | < 0.001, *** | 25 | 6.236639 | 87.06404 | 74.82431 | 99.30378 | < 0.001, *** | 71 | -1.00969 | 45.18689 | 35.68188 | 54.6919 | 0.32 |
| | 23 | 6 | 25.43339 | 97.33877 | 92.78437 | 101.8932 | < 0.001, *** | 25 | 10.39498 | 89.7122 | 81.84409 | 97.5803 | < 0.001, *** | 71 | 0.657326 | 52.17264 | 45.58212 | 58.76316 | 0.51 |
| | 24 | 6 | 56.09762 | 97.62953 | 95.55199 | 99.70707 | < 0.001, *** | 25 | 152.0594 | 98.21161 | 97.55862 | 98.8646 | < 0.001, *** | 71 | -13.577 | 14.82701 | 9.661432 | 19.99259 | < 0.001, *** |
| | 25 | 6 | 33.45513 | 97.1731 | 93.72285 | 100.6233 | < 0.001, *** | 25 | 26.18754 | 95.05773 | 91.51413 | 98.60133 | < 0.001, *** | 71 | -27.6407 | 7.315879 | 4.236734 | 10.39502 | < 0.001, *** |
| | 26 | 6 | 136.1516 | 99.33449 | 98.44786 | 100.2211 | < 0.001, *** | 25 | 51.38849 | 96.38661 | 94.52753 | 98.24568 | < 0.001, *** | 71 | -18.7939 | 11.89649 | 7.85389 | 15.9391 | < 0.001, *** |
| | 27 | 6 | 168.2234 | 99.1861 | 98.47066 | 99.90155 | < 0.001, *** | 25 | 101.0447 | 98.4893 | 97.50097 | 99.47763 | < 0.001, *** | 71 | -3.54317 | 34.88209 | 26.3744 | 43.38979 | < 0.001, *** |
| | 28 | 6 | 4.756115 | 87.09121 | 68.00863 | 106.1738 | < 0.01, ** | 25 | 10.06115 | 90.27338 | 82.02933 | 98.51743 | < 0.001, *** | 71 | -0.8802 | 46.41921 | 38.3075 | 54.53092 | 0.38 |
| | 29 | 6 | 139.4439 | 99.03938 | 98.17886 | 99.89991 | < 0.001, *** | 25 | 10.59953 | 90.28178 | 82.45485 | 98.10872 | < 0.001, *** | 71 | -2.61087 | 39.75902 | 31.93791 | 47.58014 | < 0.05, * |
| | 30 | 6 | 50.6125 | 98.09908 | 95.77369 | 100.4245 | < 0.001, *** | 25 | 15.04556 | 92.29295 | 86.5036 | 98.08229 | < 0.001, *** | 71 | -23.7594 | 11.10204 | 7.837635 | 14.36644 | < 0.001, *** |
| | 31 | 6 | 90.37333 | 98.5049 | 97.1916 | 99.8182 | < 0.001, *** | 25 | 13.68622 | 89.46632 | 83.52732 | 95.40532 | < 0.001, *** | 71 | 0.085803 | 50.31993 | 42.88511 | 57.75476 | 0.93 |
| | 32 | 6 | 3.86434 | 84.65283 | 62.71055 | 106.5951 | < 0.01, ** | 25 | 257.2368 | 99.32226 | 98.92736 | 99.71715 | < 0.001, *** | 71 | -1.73065 | 43.0251 | 34.9891 | 51.06111 | 0.09 |
| | 33 | 6 | 111.3052 | 98.88699 | 97.81227 | 99.96171 | < 0.001, *** | 25 | 80.32307 | 98.45344 | 97.21106 | 99.69582 | < 0.001, *** | 71 | -8.05963 | 22.61112 | 15.83514 | 29.38709 | < 0.001, *** |
